# *De novo* mutation of an RNA virus is increased in the presence of engineered synonymous mutations that disrupt RNA structural elements

**DOI:** 10.64898/2026.06.02.729533

**Authors:** Jeremy R. Thompson, David W. Waite, Alex Y. Cha

**Affiliations:** Plant Health and Environment Laboratory, Ministry for Primary Industries, 231 Morrin Road, St. Johns, Auckland 1072, Aotearoa New Zealand; Plant Pathology and Plant-Microbe Biology Section, School of Integrative Plant Science, Cornell University, Ithaca, NY 14853, USA

**Keywords:** Stem-loop, synonymous nucleotide change, cucumber mosaic virus, RNA, evolution

## Abstract

Using a combination of methods including selective 2’-hydroxyl acylation analyzed by primer extension sequencing (SHAPE-Seq), a complete RNA structure map of the cucumber mosaic virus (CMV) RNA 3 segment was mapped (Watters et al. (2018) Nucleic Acids Research 46, 2573–2584). To explore the effect of structural perturbations on genomic stability, infectious mutants were engineered to contain changes in one of four open reading frame (ORF) stem-loop (SL) structures SL1362, SL1439, SL1745 and SL1816, or in a highly conserved structure SL2073 in the 3’ untranslated region. The mutations occurred either in the loop or stem, thereby either maintaining or disrupting the predicted SL structure. The mutants were inoculated and then passaged on plants of *Nicotiana tabacum.* After the eighth passage, the full-length (8 kb) sequence of the virus population was sequenced by an amplicon tiling approach using the Illumina Nextera platform. All virus passages were sequenced as technical duplicates and compared against virus wild-type passages. Across the whole genome, a significant increase in *de novo* SNPs (synonymous and non-synonymous) occurred in some mutants (vs. wt) as minor (≥10%) and dominant populations, suggestive of compensatory genome-wide epistatic changes induced by changes in SLs.

**Importance:** This work further supports and expands on the novel observation that minor synonymous nucleotide changes in virus RNA structural elements can significantly increase *de novo* nucleotide changes across the whole virus genome, inducing, in some instances, amino acid substitutions. This phenomenon has broad implications for the interpretation of natural sequence variation, and more specifically for antiviral strategies.

## Introduction

RNA viruses dominate the known global virome, especially for eukaryotes, infecting animals and plants with abundant diversity (1). Of their three genomic forms; positive-sense, double-stranded RNA viruses, and negative-sense (2), the former uses the simplest replication and expression strategy with the same RNA molecule functioning as both the genome and mRNA; a property so far unique in the biological world that some have speculated its ancestors were the first replicators to appear in the primordial RNA world (3). This dual function, in a typically compact genome (4), undergoing some of the highest known mutation rates (5), has provided researchers with invaluable insights not just into virus evolution but into broader mechanisms of evolution itself (6).

Historically, genomic regions were thought to operate largely as independent units - each segment or open reading frame (ORF) associated with a predictable, linear role in replication, protein synthesis, and virion assembly. However, with ever more sophisticated tools continuing to deepen our knowledge of the mechanisms of virus evolution and the diversity of virus populations, there is growing evidence that RNA structure influences the rate and type of changes across the genome (7, 8, 9). There has also been a significant shift in the traditional view of RNA virus genomes functioning as discrete transcriptional and translational domains (10, 11, 12, 13). Instead, the emerging view is one of a dynamic multifunctional molecule that forms part of a highly interactive and evolutionarily optimized network of functional signals, one in which structural configuration, sequence plasticity, and spatial-temporal dynamics collectively shape viral replication and host adaptation (14, 15).

Cucumber mosaic virus (CMV) (species *Cucumovirus CMV;* family *Bromoviridae*) is one of the most studied of plant viruses (16) and often described as one of the most successful (17, 18) based partly on it having one of the broadest host ranges in the plant kingdom (19, 20), a property that has expanded to include a fungal host (21). It is transmitted by over 80 aphid species in a non-persistent manner (22).

In this study, we follow up on observations first made by Watters et al. (23) and later supported by Mochizuki et al. (24) that mutants with engineered synonymous changes in the coat protein (CP) ORF of CMV, when passaged *in planta*, appear to accumulate *de novo* amino acid (aa) changes in the CP at a greater rate than wild-type controls. In the latter study, when mutants carrying synonymous mutations in the 5′ half of the CP gene, were serially passaged in *Nicotiana benthamiana*, up to three *de novo* aa substitutions were observed in each CP. Similarly, Watters, Choudhary et al. (23) found fixed *de novo* distal CP aa changes when passaging mutants carrying engineered synonymous CP mutations designed to specifically disrupt structural stem-loops. Both studies speculated on the function of these *de novo* aa substitutions based on the CP’s role in virus movement (cell-to-cell and systemic). However, considering the CP’s roles in encapsidation, transmission (see (9) and references therein) and gene silencing (25), their possible function may have a broader reach within the virus cycle.

To explore this observation further, we tested whether these *de novo* aa changes were limited to just the CP or if they applied across the whole genome, evidencing a broader epistatic phenomenon. To do this, we passaged *in planta* synonymously engineered mutants, including those used by Watters et al (2018) (23), and analyzed all coding regions of the evolved virus populations by high-throughput sequencing. The results support the occurrence of these *de novo* aa changes both in the CP (confirming previous observations) and across the whole genome at varying levels of dominance, while also identifying a significant increase in the frequency of single nucleotide polymorphisms (SNPs) for some mutants, suggesting that small synonymous changes can induce a significant increase in genomic entropy.

## Materials and Methods

### Virus transcript inoculation and propagation

CMV strain Bn57 mutants were engineered as described previously, and based on RNA structural studies (23) (Fig. 1). The mutants were designed to disrupt either the formation of a stem or the sequence of a loop in defined viral RNA stem-loop elements (Fig. 2a). Plants of *Nicotiana tabacum* were kept in greenhouse conditions of 20 ± 3°C and 16 h:8 h (light: dark). Initial inoculations were carried out by rubbing 4µg of a mix of Bn57-CMV RNA1, RNA2 and RNA3 transcripts generated from infectious clones pBn57-1, pBn57-2 and pBn57-3 (or their mutant derivatives) (26), respectively, on a carborundum-dusted lower leaf of a healthy plant at a 4–6 leaf growth stage. Once symptoms appeared in the inoculated plant, approximately 100 mg of symptomatic leaf material was ground in 5 ml 0.1 M Na_2_HPO_4_ with a sterilized mortar and pestle and passaged by rubbing onto a carborundum-dusted leaf of a healthy plant at a 4–6 leaf stage of growth.

**FIG 1.**
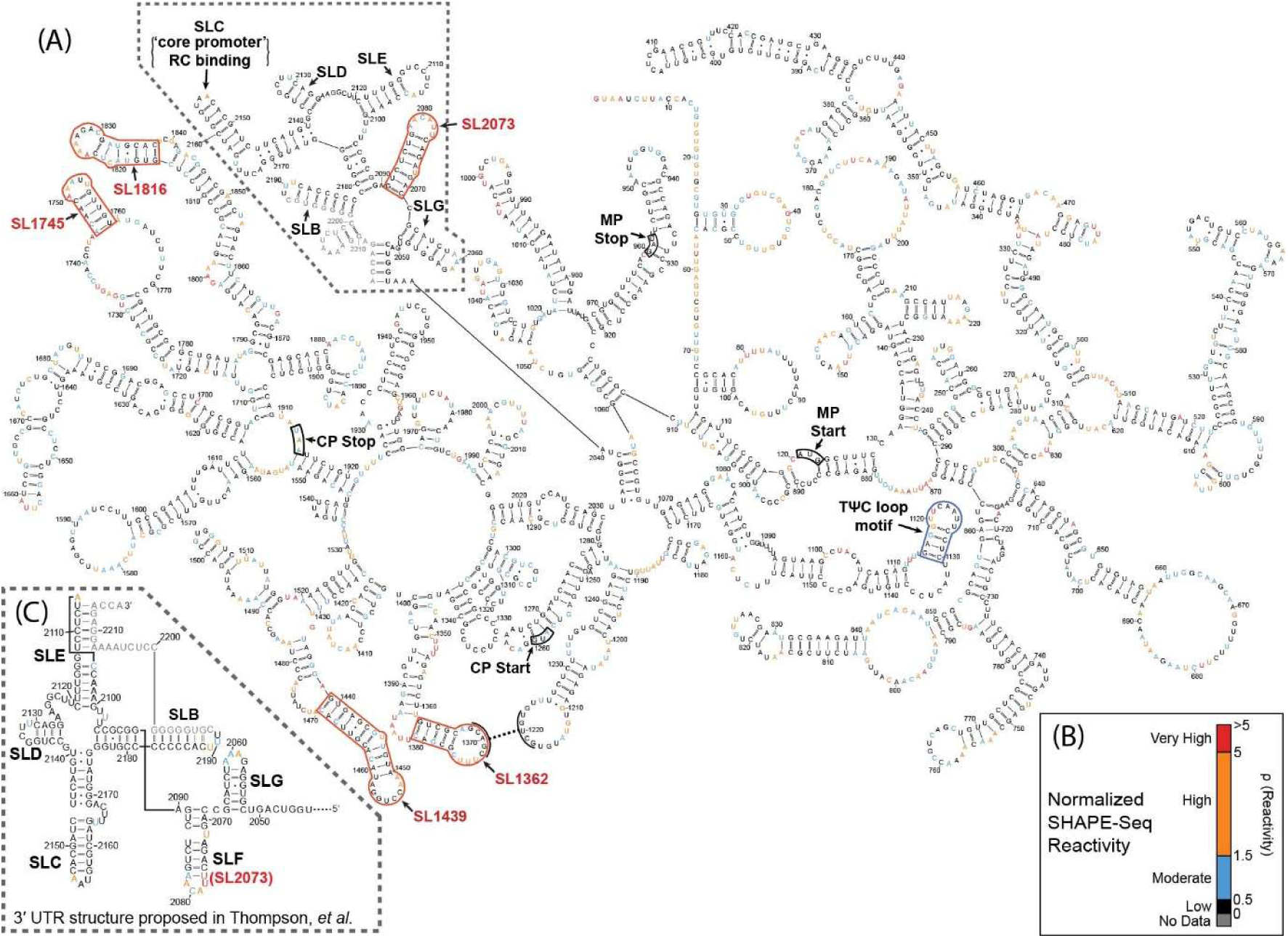
(A) Predicted secondary structure of cucumber mosaic virus (CMV) RNA3 as described by Watters et al (2018). Nucleotides are colored by the average SHAPE-Seq (Selective 2-hydroxyl acylation analyzed by primer extension sequencing) reactivity intensity as indicated in panel (B). The five stem loops (SL) analyzed in this study are outlined in red. The start and stop codons for the coat protein (CP) and movement protein (MP) are boxed in black. Alternative structures of the 3’ tRNA-like region are boxed with a dotted line, with the structure proposed by Thompson et al. (2009) in panel (C). Numbers in the sequence - nucleotide position in CMV strain Bn57 (HF572916) represents the minimum free energy structure prediction of the CMV RNA3. Original in Watters et al. Probing of RNA structures in a positive-sense RNA virus reveals selection pressures for structural elements. Nucleic Acids Research (2018) 46 (5) 2573–2584 - by permission of Oxford University Press.

**FIG 2.**
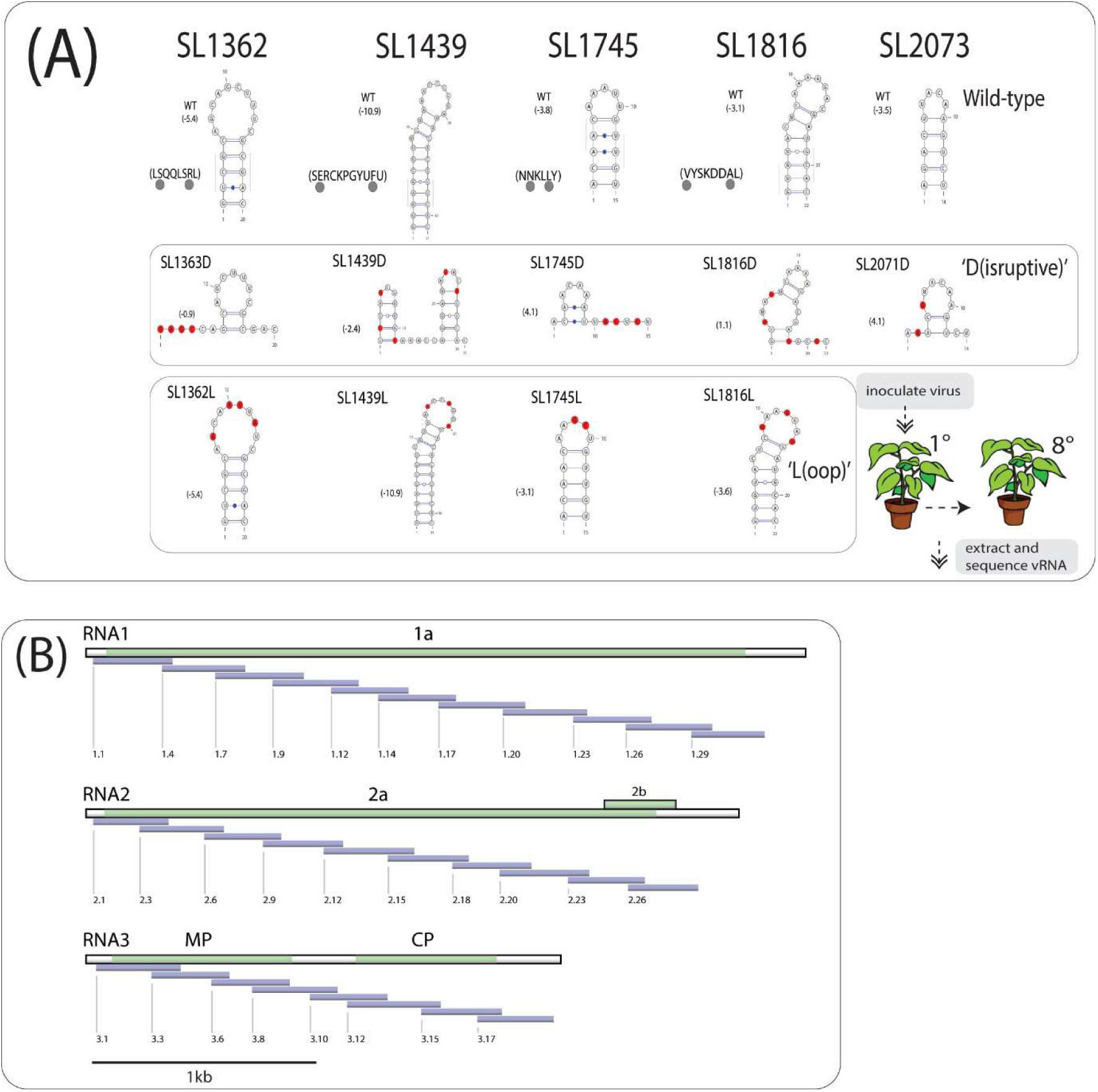
Virus passaging. (A) Schematic of engineered cucumber mosaic (CMV) mutants used in this study, with wild-type stem loop (SL) structure and free energy (number in parentheses) shown at top. Predicted amino acid sequence around SL is shown, where grey dots correspond to the codon position marked by the gray line on the SL. Predicted ‘Disruptive’ and ‘Loop’ structures are boxed, where the engineered nucleotides are shown in red. These engineered mutants were inoculated onto plants of *Nicotiana tabacum* and then passaged seven times before total nucleic acids were extracted from the eighth plant. Sequence analysis of the viral RNA was done by PCR amplification using tiled CMV-specific primers (B) (Suppl. Table 3) before being indexed for Nextera (Illumina) sequencing. (B) Position and names of the derived amplicons within the genome of CMV are shown as blue/gray bars. The boxes depict each CMV RNA segment with coding regions (green) and their names. White areas of the boxes are untranslated regions.

### RNA purification, amplicon tiling and sequencing

Tiling primers were designed using the software PCRTiler (27) to cover almost the whole viral genome, where technically practical: avoiding regions in the 5’ and 3’ untranslated region containing common sequences in all RNAs (Fig. 2b). To allow for the optimum combination of primers, amplicon sizes varied in size from 350 to 450 bases to allow for upstream pair-ended (2 x 250) Illumina (San Diego, CA) (Nextera compatible) reads. RNA was extracted from infected plant material using the RNeasy Plant Mini kit (Qiagen), and 400 ng used as a template for reverse transcription with random hexamers using M-MLV Reverse Transcriptase (ThermoFisher Scientific) according to the manufacturer’s instructions. Approximately 50 ng (0.2µl) of the cDNA generated was then amplified in two separate tubes (to avoid preferential amplification of shorter overlapping amplicons) containing two sets of tiled primers (Suppl. Table 1) by PCR using the QIAGEN® Multiplex PCR Plus kit in a total reaction volume of 12µl with the following cycling conditions: 95°C for 5 mins (x1); 95°C for 30s, 57°C for 90s, 72°C for 30s (x30); 68°C for 10 mins. Quality of the PCR products was assessed using a Fragment Analyzer (Advanced Analytical Technologies). Each of the sample’s two reactions was then pooled together, loaded into a well of a 96 well-plate, and submitted for indexing using standard Nextera primers (8bp i7 and i5 barcodes) with a MiSeq (Illumina) v2 500bp kit run and sequenced on a MiSeq (paired-end 250bp) at Cornell Biotechnology Resource Center Genomics Facility (<u>RRID:SCR_021727</u>).

All samples (n = 48) were amplified as technical replicates (wells 49 to 96 being duplicates of 1 to 48). Seven samples, inoculated with the wild-type Bn57 virus as previously described (26), were included from two separate experiments with the same growing conditions as the mutant viruses. An additional control set of amplicons derived from the DNA clones of the infectious virus sequence in plasmids (wells 37-38) was also included to detect possible repeatable *in vitro-derived* mutations during sequencing.

### Sequence analyses

Raw sequence reads were stripped of primer sequences using cutadapt v4.4 (28), then quality filtered using fastp v0.23.2 (29) with a sliding window approach (window size 4 nts, and mean quality Q20) and the resulting sequences inspected using FastQC to verify removal of primers and barcodes. Filtered reads were then mapped to the CMV strain Bn57 reference genome (NCBI sequence accessions HF572914.1, HF572915.1, and HF572916.1) using bowtie2 v2.5.0 (30) with the --very-sensitive mapping profile. Unaligned reads were removed from each resulting sam file, and reads were sorted and compressed in bam format using samtools v1.16.1 (31).

Mapping files were summarised using the pysam library v0.20.0 (37), tallying the number of A, T, C, G, insertions, and deletions at each position of the reference Bn57 genome. We reported only mutations at positions with a mapping depth of at least 100-fold, and where the SNP was present in at least 10% of the reads mapping to the position. To eliminate the chances of PCR or sequencing error giving rise to false positive SNPs, we then filtered the SNPs to retain only mutations meeting the above criteria and that were also shared between technical replicates. SNP abundance profiles were visualised using the plotly v5.13.0 library in python (32).

The filtered SNPs were then assessed in the context of their codon within the canonical coding regions annotated in the reference Bn57 genome. Each SNP falling within the coding region was characterised in terms of its impact on protein translation, either as a synonymous or non-synonymous mutation. To test for the prevalence of non-synonymous mutations arising during this work in the global population, CMV sequences were downloaded from NCBI GenBank using the datasets command line tool (date of download: 21-Mar-2025). Each sequence was aligned to the Bn57 genome using the Geneious reference mapping tool with the ‘High Sensitivity’ mapping profile.

### General Linear Model (GLM) and rate-ratio estimation

Per-individual substitution rates were compared between mutant and wild-type groups using a Poisson GLM with a log link. Total substitution counts for each gene and group were entered as the response in a Poisson GLM, with the number of individuals contributing to each total included as a log-offset so that the model estimated per-individual substitution rate differences between mutants and wild-type to account for unequal sampling. The model was:

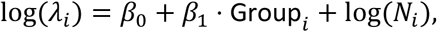

where *N_i_* is the number of individuals contributing to each count. The exponentiated group coefficient, exp(*β*_1_), was interpreted as the mutation rate ratio (mutant/wild-type). Significance was assessed using a Wald test based on the ratio of the coefficient to its standard error.

## Results

### *De novo* single nucleotide polymorphisms occur at a significantly higher frequency in some stem-loop mutants than in wild-type virus

We restricted our analysis to only SNPs shared between both technical replicates, as these SNPs would most likely be *in vivo* in origin and therefore biologically relevant. We then looked at SNPs present in ≥10% of sequences in the population rather than fully fixed mutations in order to capture emerging within-host heterozygosity (Fig. 3). A cut-off ≥10% was chosen for ease of data analysis: cut-offs of 10%, 5% and 2.5% showed consistent significant differences using the Wilcoxon Signed-Rank test between mutants (SL1362L, SL1362D, SL1745L and SL2073D) and wild-type (Suppl. Fig. 1). There was no significance detected with cut-offs of 25% and 50%. Averaging the total number of SNPs (≥10%) for each mutant (Suppl. Table 2), we found that after passaging four mutants (1362L, 1362D, 1745L and 2073D) accumulated SNPs at a significantly higher level (average ≥25) than the wild-type virus (average 14) (P = 0.004, 0.019, 0.030, 0.017) while both mutant types (L and D) of the stem-loop 1439 resulted in notably elevated numbers of SNPs (average ≥22) in comparison with the wild-type (Fig 3). The best characterized of all the structural elements analyzed here is 2073D, which is a component of the highly conserved 3’UTR (aligning with earlier studies, 2073D is also known as SLF (33)) (Fig 1). Watters et al (23) showed that disruption of this stem-loop structure consistently resulted in a close-to-wildtype structural reversion when passaged *in planta,* while changes in its loop sequence were tolerated. Interestingly, the mutant with the highest average SNPs (n = 30) is 1362L. The loop of this stem-loop has been proposed to form a kissing interaction with a loop (UGUCG) in a predicted RNA3 intergenic stem-loop beginning at position 1205. Canonical base-pairing would suggest that the engineered sequence in 1362L would change the proposed interacting nts 1368-1372 from GCAGC to ACAAU, making only the third A in the latter non-pairing, which in all five passages of SL1362L reverted to the wild-type G, thereby supporting the proposed kissing-loop interaction. Mutation of the loop in stem-loop 1745 also had a more significant effect than disrupting the stem itself, suggestive perhaps too of a putative unidentified interaction (undetected in our dataset).

**FIG 3.**
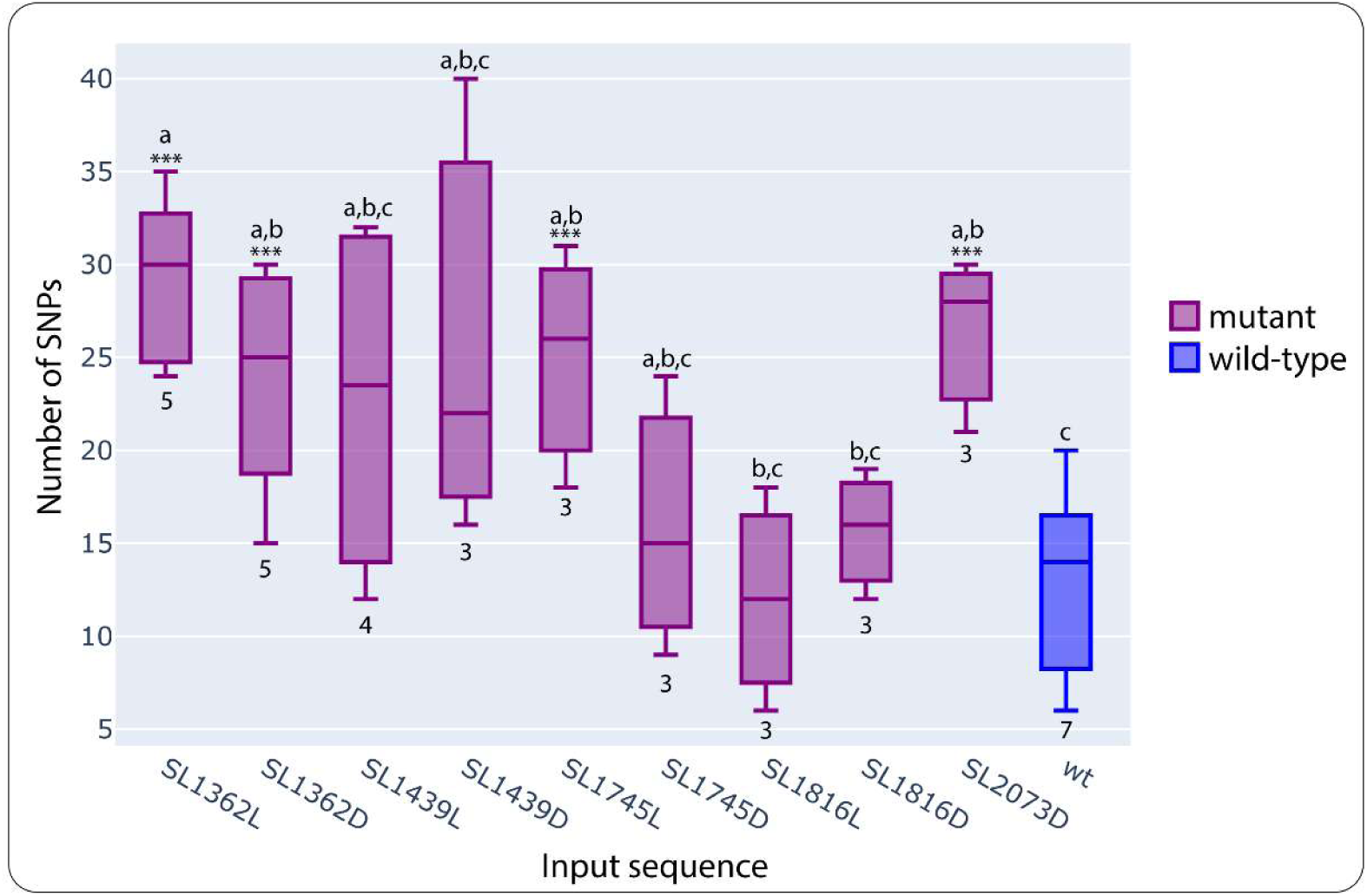
Graph showing the number of *de novo* single nucleotide polymorphisms (SNPs) for each stem-loop mutant and wild-type (input sequence) at or above a frequency of 10%. Spread of SNPs for each individual passage - vertical boxes with standard error bars. Numbers below boxes – number of biological replicates. Lettering above boxes – statistical groupings calculated by the Wilcoxon Signed Rank test, where a triple asterisk indicates a significant difference to the wild-type.

### *De novo* amino acid substitutions predominate in stem-loop mutants and occur across the viral genome

Mutants show a significant increase in *de novo* aa substitutions, both at the individual level and consistently across genes (Fig 4). When the number of substitutions was normalized by the number of individuals per group, mutants showed approximately a 1.8-fold increase in per-individual substitution rate relative to wild-type, and this difference was statistically significant (P = 0.0014; 95% CI 1.25-2.50; coefficient β = 0.569) when analyzed by generalized linear models with a Poisson regression model (dispersion value 1.18) with a log-offset for sample size. To assess whether substitution rates were consistently elevated across genes, they were treated as matched units. For each gene, substitution rates per individual in each group and derived gene-specific rate ratios were calculated as significant (P = 0.04). At the gene level, four of the five genes displayed higher substitution rates in the mutants (including the 2b gene, which had no substitutions in the wild-type) (Table 1). Notably, the replication-associated genes (1a and 2a) had the highest ratios (2.14 and 4.14), while the MP was the only gene for which the wild-type had more substitutions.

**FIG 4.**
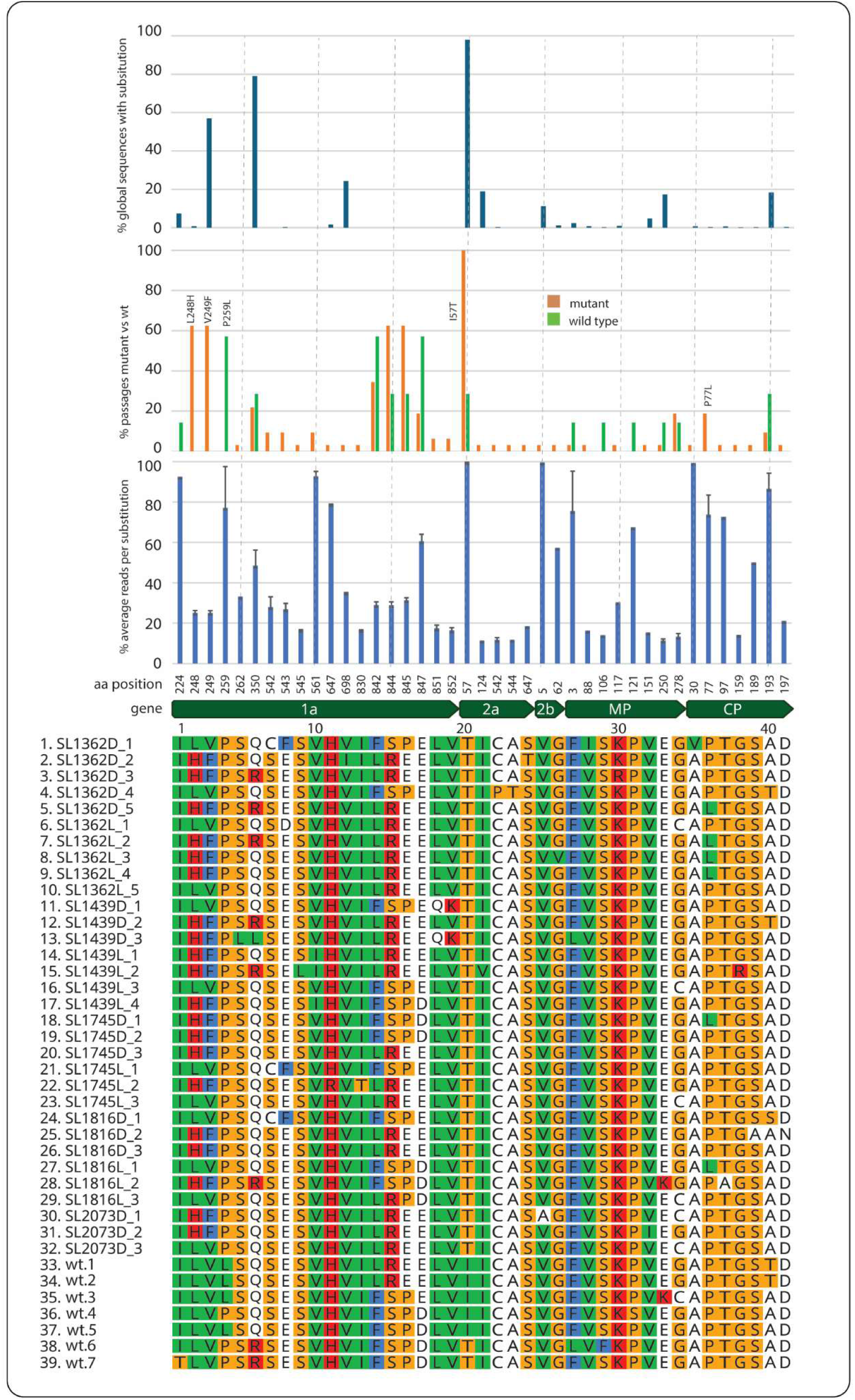
Predicted *de novo* amino acid substitutions (≥10%) for each passaged virus (1-39 lower left) across the entire proteome of cucumber mosaic virus (CMV). Stem-loop (SL) mutants are numbered as described in Figures 1 and 2; the letters ‘D’ and ‘L’ indicate a ‘Disruptive’ or ‘Loop’ mutant, respectively. The alignment shows only those amino acids (n = 41) that underwent substitution. The alignment is colored according to the Clustal X scheme (Geneious, Biomatters). The position of each residue is shown, first, with reference to the virus coding region (gene) with each open reading frame labelled within the dark green arrow, and second, by its amino acid position (vertical number) within the gene’s open reading frame. All histogram bars are aligned with their corresponding amino acids shown in the alignment. The lower histogram shows the average number (with standard error bars) of reads containing each substitution as a percentage of the total number of Illumina reads. The middle histogram shows the percentage of individual passages (mutant (n = 32) vs wild-type (n = 7)) containing the identified substitution. The upper histogram shows the percentage of global (GenBank) CMV sequences that contain the substituted amino acid recorded here.

**Table 1.** *De novo* amino acid substitution rates in cucumber mosaic virus mutants and wild-type for each viral gene. Rates capture the average number of substitutions per individual. A Haldane-Anscombe correction (0.5 added to each mutant and wild-type count) was applied to avoid an undefined rate ratio for the 2b gene, which had zero wild-type substitutions.

| Gene | Mut no. | WT no. | Mut rate | WT rate | Rate ratio |
| --- | --- | --- | --- | --- | --- |
| 1a | 186.5 | 19.5 | 5.83 | 2.79 | **2.09** |
| 2a | 38.5 | 2.5 | 1.20 | 0.36 | **3.33** |
| 2b | 2.5 | 0.5 | 0.08 | 0.07 | **1.10** |
| MP | 43.5 | 12.5 | 1.36 | 1.79 | **0.79** |
| CP | 14.5 | 2.5 | 0.45 | 0.36 | **1.25** |

The most conspicuous differences in aa substitution between the mutants and wild-type occurred in the 1a protein at residues L248H, V249F and P259L. Substitutions L248H and V249F occurred at a population frequency of at least 25% and always together in 20 derived mutants. Sixteen of these appeared to be associated with distal 1a substitutions at positions S844R and the substitution P845E, which are predicted to have significant effects on protein stability, structure and function. Even more disruptive is the substitution at residue P259L, which occurs exclusively in 4 of the 7 mutants at a population frequency of 77% (Fig. 4).

The *de novo* substitution at residue P77L in the CP, identified by Watters et al (23), appeared in 3 of the 4 SL1362L derivatives and also in one derivative each of SL1362L, SL1745D and SL1816L at varying levels of fixation within the population (40-100%). Examples of fully (or close to) fixed substitutions in derivatives are limited to three examples: I57T in the 2a, V5A in the 2b and A30V in the CP. V5A and A30V are single events in passages SL162D1 and SL2073D1, respectively, whereas I57T occurs in 34 of the 39 passages, including two wild-type passages. It also occurs in 98% of global CMV sequences (n = 473) (Fig 4) (Suppl. Table 3), suggesting a strong natural selection. Amino acids, also prevalent in CMV global sequences and conspicuous in the passaged population here, are *de novo* substitutions Q350R (1a) represented by 79% (n = 607), and the unique-to-mutant V249F (1a) represented by 58% (n = 654).

Interestingly, 2a substitutions, F842L, S844R, P845E and D847E do not register at all in global CMV sequences but are quite dominant in these passaged viruses (both mutant and wild-type), suggesting they might occur consistently as subpopulations and are therefore never recorded in GenBank or that they are unique to this strain of CMV.

## Discussion

The results presented here demonstrate that even a small number of synonymous perturbations to local RNA secondary structures can have significantly broad effects on the virus genome. By engineering targeted synonymous mutations into five stem-loop (SL) elements of CMV RNA3 and serially passaging these viruses *in planta*, we observed a consistent and significant increase (1.8x) in *de novo* SNPs relative to wild-type controls. This finding confirms and expands on earlier observations that synonymous mutations in the CP ORF can result in proximal amino acid substitutions (23, 24) and shows that the phenomenon is not restricted to the CP but instead reflects a genome-wide epistatic response. The fact that both loop-altering and stem-disrupting (SL1362L, SL1362D, SL1745L, and SL2073D) engineered mutations produced elevated *de novo* mutation rates indicates that the virus is sensitive not only to the loss of local base-pairing but also to changes in putative tertiary interactions such as kissing-loop contacts.

A notable outcome of this study is the concentration of observed *de novo* amino acid substitutions in the replication proteins 1a and 2a, suggesting that these proteins are significantly affected by RNA structural perturbations elsewhere in the genome. This implies that the replicase complex may be a primary target for compensatory adaptation, potentially adjusting its fidelity, processivity, or interaction with structured RNA elements.

Consistent with earlier studies, the CP substitution P77L, repeatedly observed across multiple mutants, maps to between βB and βC strands within the CP shell domain (34, 35). Replacing proline with a leucine would possibly stabilize the β-strand and subtly alter capsid rigidity. This may compensate for changes in RNA folding or packaging efficiency induced by the engineered synonymous mutations. Another explanation is that the aa change is driven by underlying RNA structural requirements. However, the aa change is caused by the mutation of a cytosine for uracil at nt position 1487, located in a region predicted to be unpaired and unstructured (23). The presence of I57T in the CP of 34 of 39 passages, including wild-type controls, and its prevalence in 98% of global CMV sequences, provides a potential example of evolutionary convergence. It is therefore important to highlight that these genome-wide non-synonymous changes are not limited to mutants, only that their mutation rate across the genome is elevated.

Taken together, these findings support the idea that disruption of a single stem-loop can propagate selective pressures across the entire genome, altering mutation rates and driving amino-acid changes in distant proteins. This aligns with emerging views that RNA structure acts as a “puppet master” of viral evolution, shaping not only local translation and replication but also long-range interactions and genome-wide mutational landscapes (9).

Our data also highlights the importance of considering synonymous mutations as potentially impactful evolutionary events. Although traditionally viewed as neutral, synonymous changes that alter RNA structure can induce substantial fitness costs (36), and in our study, reshape the viral proteome. Protein misfolding has been shown to occur due to changes in codon translation rates induced by synonymous mutations (37, 36), but we have been unable to find any evidence in the literature of distal aa changes occurring at a higher rate due to synonymous changes. This finding may be due to our study targeting subpopulations that have been overlooked in earlier studies. Whether these subpopulations would remain at their current frequency, go extinct or become fixed would require further investigation and may differ for each combination of mutations. Nevertheless, this observation has broad implications not only more generally for the interpretation of natural sequence variation, but also more specifically for antiviral strategies (38).

While our study provides evidence of genome-wide epistatic responses to structural perturbation, it is important to note that our sequencing approach captures consensus-level and dominant subpopulations of both genomic and messenger RNA variants but does not resolve full haplotypes; long-read sequencing could clarify whether specific combinations of mutations co-segregate.

In conclusion, this work provides strong evidence that minor synonymous mutations disrupting RNA secondary structure can induce significant changes across the entire CMV genome. The resulting increase in genomic entropy uncovers an evolutionary connection between RNA structural elements and protein-coding regions in positive-sense RNA viruses.

## Data availability

All data supporting the conclusions of the study are included within this article and the BioProject number PRJNA1465633.

## Acknowledgements

We would like to thank Peter Schweitzer, James Vanee and Linda Cote at the Cornell Genomics Facility (<u>RRID:SCR_021727</u>) for their technical support and sound advice. A big thank you also goes to Keith

Perry, Heather McLane, Chad Thomas and the greenhouse crew for their support and benevolence. This work was in part funded by the Cornell Institute of Biotechnology Seed Grant program (U353710)

## Supplementary data

Suppl. Fig. 1 – SNP frequencies at different screening depths.

Suppl. Table 1 – Tiling primers used for the sequencing of cucumber mosaic virus

Suppl. Table 2 – Table of all SNPs and aa substitutions

Suppl. Table 3 – Analysis of global sequences containing study-specific substitutions.

